# Two divergent haploid nuclei shaped the landscape of population diversity in wheat stripe rust

**DOI:** 10.1101/2024.12.10.627719

**Authors:** Yibo Wang, Mou Yin, Fei He

## Abstract

Heterozygosity is a measure of allelic diversity within individuals. *Puccinia striiformis* f. sp. *tritici* (*Pst*) is a highly heterozygous dikaryotic crop pathogen. Its source of heterozygosity variation and the contribution to adaptability are still unknown. By analyzing resequencing data of 266 worldwide *Pst* isolates, we found that *Pst* may have undergone an important historical hybridization event, introducing substantial diversity and leading to the divergence of its haploid genomes into two distinct haplotypes. Strains with both haplotypes exhibit higher individual allele diversity and wider geographical distribution. Between the two haploid genomes, 16% of the genome had diverged, scattered as mosaic blocks within the genome. These regions are enriched with genes displaying critical roles during infection of the plant host, and exhibit higher expression levels. In these regions, 8.0 Mb shows recombination fingerprints associated with virulence, while 4.9 Mb displays linkage across the entire genome. We demonstrated that sexual recombination in *Pst* is extensive and significant. *Pst* can gain genetic diversity and adaptability due to intra- and inter-species hybridization. Our study resolved the debate over the sources of individual allele diversity in *Pst* and expands the understanding of pathogen virulence evolution. These findings also suggest that interrupting the sexual reproduction of pathogens can be an effective strategy for controlling wheat stripe rust.

## Introduction

*Puccinia striiformis* f. sp. *tritici* (*Pst*) is a dikaryotic rust fungus that severely threatens wheat production. It has an alternation of generations: the asexual phase infects cereal hosts, while the sexual phase lives on alternate hosts such as barberry (Berberis spp.) and Mahonia spp. During its life cycle, it produces up to five types of spores at different phases^1^. In addition to asexual and sexual reproduction, somatic hybridization was found to contribute to genetic diversity in rust fungi. For example, the Ug99 lineage of *Puccinia graminis* f. sp. *tritici* (*Pgt*), which emerged in 1998, shares a single nuclear haplotype with the older South African *Pgt21* lineage, indicating it may be an isolate of somatic hybridization^2^. Similarly, shared nuclear haplotypes among Australian *Puccinia triticina* (*Pt*) lineages 19ACT06, 19NSW04, and 20QLD87 confirm the occurrence of somatic hybridization in nature^3^.

The Meselson effect refers to a genomic characteristic of species that reproduce asexually for extended periods, with one of the most obvious features being the accumulation of heterozygosity. Heterozygosity is considered as a measure of allelic diversity within individuals and the indicator of asexual reproduction^4,5^. Previous studies used limited molecular markers to assess the heterozygosity in *Pst*, revealing high heterozygosity and population differentiation within the species based on heterozygosity levels^6,7^. In regions like the Himalayan and Africa, the lineages of *Pst* have low heterozygosity and no deviation from Hardy-Weinberg equilibrium (HWE). However, lineages in Europe, the Americas, and Oceania exhibit high heterozygosity and deviation from HWE^7^. The explanation of this phenomenon remains controversial among scholars. Some suggested that the high-heterozygosity lineages are products of the Meselson effect. They undergone long-term asexual reproduction, while low-heterozygosity lineages may have experienced recent sexual reproduction events^7-20^. A recent study proposed that high-heterozygosity lineages are the progeny of a somatic hybridization event between two highly differentiated lineages, rising question to origin of the genome heterozygosity in this serious wheat pathogen^18^.

In fact, debates concerning heterozygosity similarly emerged in studies of bdelloid rotifers. Initially regarded as organisms likely reproducing exclusively through asexual reproduction, bdelloid rotifers were found to exhibit high levels of heterozygosity in the past century. However, with the advent of more sequencing data, scholars discovered that heterozygosity in bdelloid rotifers originated from paralogous genes. The genome of bdelloid rotifers is enriched with exogenous genes. These exogenous genes contribute to the increased heterozygosity^21,22^.

We still do not clearly understand how the complex life cycle and alternative reproductive strategy impact the evolution of *Pst*. The heterozygosity population divergence (hpd) is particularly intriguing. It may play an important role in understanding the virulence evolution. Notably, several recent studies provided extensive *Pst* sequencing data, including hundreds of field isolates and high-quality haplotype-level genomes. Massive public data offer opportunities for in-depth studies of *Pst* population diversity and hpd^12,23-37^. Here, we developed a single-nucleotide polymorphism (SNP) calling pipeline, Dfseq-calling, and applied it to detect diversity in sequenced isolates worldwide. This analysis revealed the driving forces of hpd and the impact of hpd on the diversity and adaptability landscape, highlighting the global A and B haplotype differentiation and its evolutionary significance on this wheat pathogen.

## Results

### Construction of the global *Pst* genetic variation map using Dfseq-calling pipeline

We first gathered 266 whole genome sequencing (WGS) public data of worldwide *Pst* isolates (Supplementary Table 1). After preliminary quality control filtering, reads were aligned to the reference genome Pst134pri^23^ for SNP calling. However, after applying stringent filtering steps, the resulting genetic variation map still exhibited unexplained errors. At the individual level, some isolates displayed linkage lossing, with large regions showing a mosaic distribution of genotypes (Supplementary Fig. 1a). At the population level, some regions exhibited a mosaic genotype distribution across all isolates (Supplementary Fig. 1b). The failure of the standard SNP calling pipeline to perform high-quality variant detection on public datasets suggests that some samples in these datasets may be of poor quality. After inspection of the data and pipeline, we identified the causes of these errors.

At the individual level, allele balance (AD) values were calculated for heterozygous SNP sites in each isolate with mosaic genotypes. Its AD values showed an abnormal bimodal distribution instead of the normal distribution around 0.5 observed in regular isolates. This indicates potential strain mixing events (Supplementary Fig. 1a). At the population level, regions with mosaic genotypes enriched with transposable elements (TEs) and lacked of genes. A focused analysis of TE regions showed a significant mosaic genotype pattern, likely caused by multiple highly similar reads aligning to the same reference genome position, leading to SNP calling errors (Supplementary Fig. 1b).

To avoid these issues, we improved the SNP calling pipeline and developed Dfseq-calling, a workflow tailored for variant detection in dikaryotic microorganisms (Fig. 1). This pipeline effectively differentiates true heterozygous SNP sites from erroneous sites caused by strain mixing (Supplementary Fig. 1c), in order to do more accurate downstream analyses.

**Fig.1.**
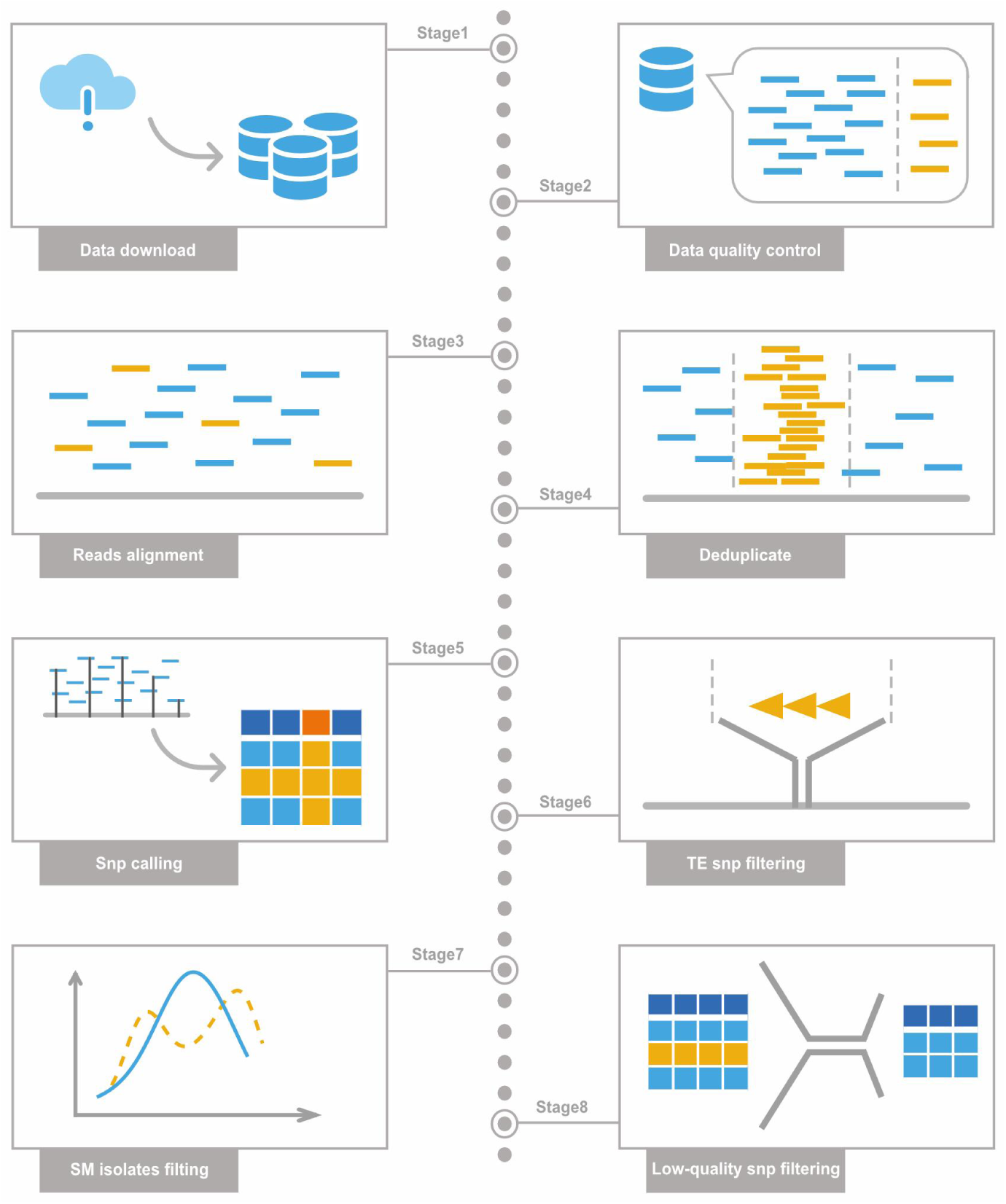
Dfseq-calling pipeline. Using the Dfseq-calling pipeline for detecting genetic variation in dikaryotic microorganisms effectively eliminates risks of strain-mixing. First, 266 public WGS data were downloaded from NCBI and subjected to quality control. These reads were aligned to a reference genome, generating BAM files. BAM files were then deduplicated. SNP calling was performed, during which TE regions causing errors were masked. Strain-mixing risks were evaluated by assessing allele balance of heterozygous SNPs. Isolates with strain-mixing risk were removed. Finally, filtering of low-quality variants produced a cleaner VCF.

Using the Dfseq-calling pipeline, we removed 27 potential strain-mixing isolates and masked TE sequences in the reference genome. A total of 8.98 billion reads were generated from 239 *Pst* WGS public data, with an average of 37.6 million reads per isolate. On average, about 83.9% of the reads were uniquely mapped to the Pst134pri genome, with an average coverage depth of 45.5× (Supplementary Table 2). Finally, we constructed a high-quality global *Pst* genetic variation map. This map contains 1,008,364 SNPs for a population of 239 isolates published by 14 studies (Supplementary Table 3). The regions between 3-4 Mb on chr9 and 1.4-2.1 Mb on chr17 show lower SNP density (Supplementary Fig. 2a). Interestingly, the region from 2.8 to 4.2 Mb on chr9 was reported to undergo genome degeneration, even a lack of genes. The *Pr* gene controlling mating type is located at the start of this region (2.98-2.99 Mb)^38^. When calculating the minor allele frequency (MAF) and missing rate of each SNP, a bimodal distribution was observed, with many SNPs having a MAF around 0.2 (Supplementary Fig. 2b), and some SNPs showing F_MISS around 0.36 (Supplementary Fig. 2c). We used Fis (fixation index, Fis) to represent the heterozygosity of each isolate. Consistent with previous studies, we found the isolates were categorized as high and low heterozygosity (Supplementary Fig. 2d). The re-discover of these patterns here in our study suggested our strict SNP calling procedure is necessary to make use of public WGS data.

### Genetic diversity and heterozygosity of *Pst*

The 239 isolates were collected from six continents and 15 countries, with most isolates concentrated in Asia, Africa, Europe, and North America, while less isolates were collected from South America and Oceania (Fig. 2a). To understand the evolution and phylogenetic of *Pst*, we constructed a phylogenetic tree (unrooted) using SNPs from the gene regions of the genetic variation map (Fig. 2b). We found that the differentiation in heterozygosity within global isolates was more pronounced than geographic differentiation, with high- and low-heterozygosity isolates clustering into two branches. A small number of low-heterozygosity samples from Asia and Africa were grouped into the high-heterozygosity branch. Geographically, isolates from same continent clustered together, but isolates from Asia and North America were split into multiple branches. For example, Asian isolates were divided into two branches, including isolates with high and low heterozygosity, respectively. Notably, occasional isolates from other continents were intermixed within extraneous branches, suggesting intercontinental strain spread events.

**Fig.2.**
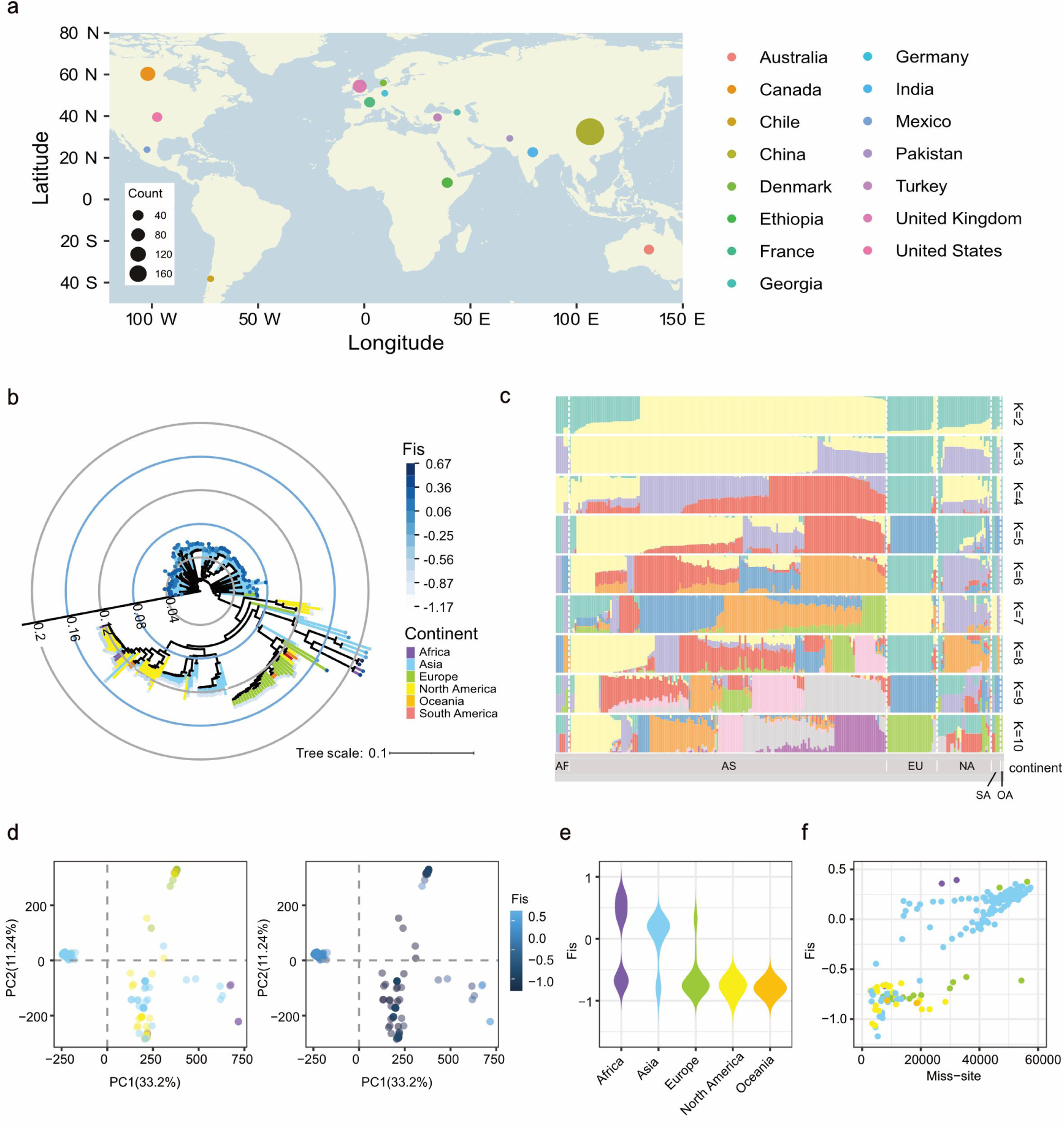
Population diversity and hpd of *Pst.* a. Geographic distribution of 239 worldwide *Pst* isolates. Different colors represent sampling countries, and circle size indicates the number of isolates. b. Phylogenetic tree of worldwide *Pst* isolates constructed using SNPs in gene regions. Branch colors represent the sampling continent of isolates and leaf node colors indicate their Fis. c. Ancestral components of global *Pst* isolates. The x-axis represents the sampling continent of isolates and each color denotes an ancestral component. d. PCA of global *Pst* isolates, with the left panel’s point color showing isolates sampling continent and the right panel showing Fis. The first two components explain a large portion of genetic variation. e. Distribution of isolates’ Fis from different continents. The x-axis and violin plot colors represent the sampling continents and the y-axis shows heterozygosity. f. Correlation between missing loci and Fis in isolates, where the x-axis shows the number of missing sites, the y-axis shows Fis, and the colors represent the sampling continent.

We estimated the ancestral component for each isolate (Fig. 2c). We found that most European isolates shared a single ancestral component. A few recent sampling isolates exhibited different ancestry, possibly indicating the spread of Asian strains into Europe. Asian isolates displayed multiple ancestral components, suggesting that this region is the origin of diversity. Principal component analysis (PCA) also indicated significant hpd (Fig. 2d). Interestingly, within the low-heterozygosity group, isolates were divided into two clusters, while most high-heterozygosity European isolates formed a distinct cluster. The diversity and phylogenetic appear to be related to heterozygosity in *Pst*. Low-heterozygosity isolates were mainly distributed in Asia and Africa, while high-heterozygosity isolates were found in Europe, North America, and Oceania (Fig. 2e). These findings are consistent with previous studies suggesting that the Himalayan region of Asia and the Middle East are the centers of diversity for *Pst.* In these regions, frequent sexual reproduction may have occurred, resulting in the emergence of low-heterozygosity isolates. In contrast, isolates from other continents are more likely to be products of the Meselson effect. They undergone long-term asexual reproduction due to the absence of alternate hosts^7^.

However, the phylogenetic tree indicated that the high-heterozygosity isolates are significantly related. It is hard to explain why they did not diverge as a result of long-term asexual reproduction. When counting the SNP missing rates for each isolate, we unexpectedly found that it also aligned with hpd: high-heterozygosity isolates had lower SNP missing rates, while low-heterozygosity isolates had higher SNP missing rates (Fig. 2f). The unusual distribution of various indices, such as MAF and F_MISS (Supplementary Fig. 2b, c), may be related to hpd in *Pst*. All these suggesting the necessity for deeper exploration of hpd. Also, this raises questions about previous conclusions. Is it sexual or asexual reproduction that drives hpd in *Pst*? Does the Meselson effect drive high heterozygosity in *Pst*? Overall, our analyses highlight that understanding how the cause of heterozygosity is crucial for comprehending the evolution and diversity of *Pst*.

### A/B haplotype combinations result in hpd

Previous studies found different combinations of a limited number of haploid genomes in Pt^3^, indicating that *Pst*, another wheat rust pathogen, may also possess distinct nuclear combinations. We compared the four nuclei of two previously published high-quality haplotype-level genomes, AZ2 and Pst134^23,24^. Pairwise SNP variation detection of the four haploid genomes (Fig. 3a) revealed that AZ2 from China is a low-heterozygosity isolate, while Pst134 from Australia is a high-heterozygosity isolate. Interestingly, haplotype comparison between the two isolates also revealed a heterozygosity difference. Fewer SNPs were detected when comparing both AZ2A-Pst134alt and AZ2B-Pst134alt, while more SNPs were detected when comparing AZ2A-Pst134pri and AZ2B-Pst134pri. Based on this, we hypothesized that the two haploid nuclei in AZ2 and Pst134alt are more closely related and may originate from same ancestor, which we define as A haplotype. As a contrast, Pst134pri is defined as B haplotype. Collinearity analysis supported this hypothesis (Fig. 3b and Supplementary Fig. 3), with A-A comparisons showing higher alignment identity and less noise, while A-B comparisons exhibited lower identity and more noise.

**Fig.3.**
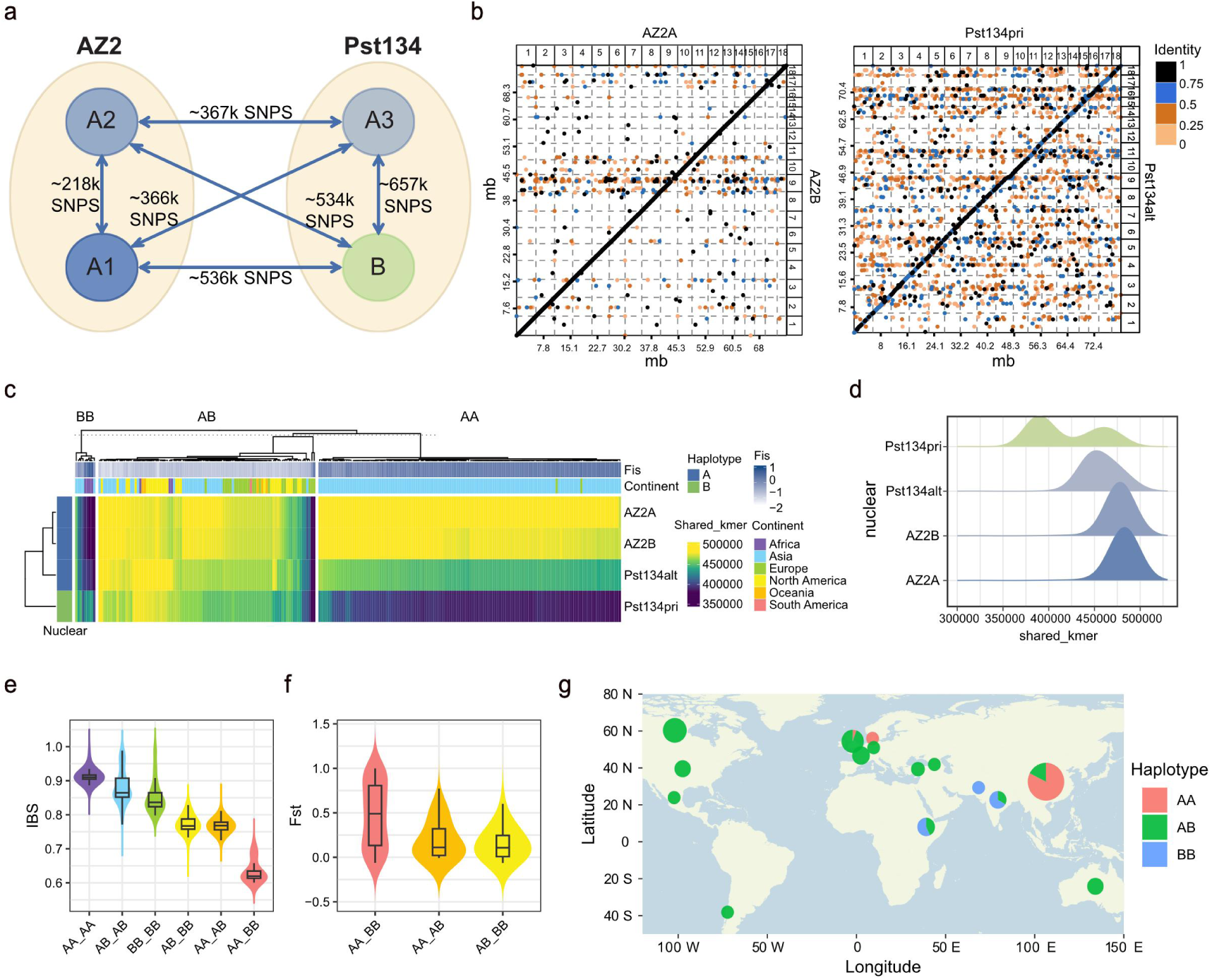
Reason for hpd: A/B haplotype hypothesis supported by multiple results. a. SNP variant counts between four high-quality haploid genomes. The numbers next to the arrows indicate the SNP counts. b. Collinearity analysis for different haplotypes. Point and line colors indicate sequence identity. AZ2A, AZ2B, and Pst134alt represent A haplotype, while Pst134pri represents B haplotype. c. A/B haplotype combinations of global *Pst* isolates. The heatmap’s x-axis represents the haploid genome (four rows for four distinct genomes) and each column represents an isolate. The heatmap colors show the shared k-mer count between the isolates reads and the haploid genome. Top annotation shows Fis and geographical origin, while left annotation indicates the haplotype (A/B) of the four haploid genome. The clustering reveals three major haplotype combinations. Low-heterozygosity BB haplotype isolates share fewer k-mers with A haplotype genome and more with B haplotype genome. Low-heterozygosity AA haplotype isolates share fewer k-mers with B haplotype genome and more with A haplotype genome. High-heterozygosity AB haplotype isolates share more k-mers with both A and B haplotype genomes. d. Distribution of the shared k-mer count between global *Pst* isolates and the four haploid genomes. e. IBS among isolates with different haplotype combinations. f. Population genetic divergence (measured by *F*_ST_) among subgroups of isolates with different haplotype combinations. g. Geographic distribution of isolates with different haplotype combinations.

After defining the haplotypes A and B, we aimed to extend this identification into population level in order to test if the divergence between the two haplotypes is the reason for hpd. By calculating shared-kmer counts between the WGS reads of 239 *Pst* isolates and the A/B haploid genome, we assessed the genomic distance and assigned haplotypes to each isolate. We found on a broad scale, the genome composition of most *Pst* isolates could be explained by the combinations of A and B haplotype, including AA, AB, and BB (Fig. 3c and Supplementary Table 4). AA haplotype isolates have higher k-mer sharing with A haploid genome, while BB haplotype isolates share more k-mers with B haploid genome. AB isolates share higher k-mer counts with both A and B nuclei. Only 3 isolates showed low k-mer sharing with both A and B haploid genomes, possibly due to poor sequencing quality or the emergence of a new haplotype.

The combinations of haplotypes A and B were directly linked to heterozygosity: AB haplotype isolates exhibited higher heterozygosity, while AA and BB haplotype isolates showed lower heterozygosity. By plotting the distribution of k-mer sharing between the 239 isolates and A/B haploid genomes (Fig. 3d), we observed a bimodal distribution for B-nucleus sharing, with some isolates sharing fewer k-mers with B, and more k-mers with A, likely due to the underrepresentation of BB haplotype isolates in the population. Genotype heatmaps (Supplementary Fig. 5c) also provided evidence supporting the A/B haplotypes hypothesis. There are numerous regions exhibiting heterozygosity in the AB haplotype isolates, alternative allelic homozygosity in the AA haplotype isolates, and reference allelic homozygosity in the BB haplotype isolates (Supplementary Fig. 5c).

We calculated the IBS (identity by state) for different haplotype combinations in the population (Fig. 3e). We found that isolates of the same haplotype have higher IBS identity, while those with only one different haplotype show intermediate IBS identity, and isolates with both haplotypes different have the lowest identity. *F*_ST_ (fixation index, *F*_ST_) analysis also indicated significant population divergence between AA and BB groups (Fig. 3f). The geographic distribution of haplotypes showed that AB haplotype isolates were widespread, AA haplotype isolates were concentrated in Asia, and BB haplotype isolates were mainly found in the Middle East, Africa, and India (Fig. 3g).

These findings strongly supported the A/B haplotypes hypothesis and challenged previous understandings of heterozygosity in *Pst*^7-20^. The conclusion that high-heterozygosity strains in *Pst* (AB-type) result from Meselson effect is not entirely correct. For high-heterozygosity strains across continents with no gene flow, these strains share many similar genotypes. This is difficult to explain, as they did not accumulate differentiation between each other through long-term asexual reproduction. AB haplotype isolates are more likely to be a fusion of A haplotype and B haplotype, suggesting hpd is caused by different haplotype combinations.

### Genome regions with distinct population variation landscapes

After explaining the cause of hpd, we aimed to explore which specific variations contributed to the divergence of A and B haplotypes. The genotype heatmap showed different types of population variation landscapes across the *Pst* genome (Fig. 4a). The first type of genomic region showed no significant heterozygosity differences between isolates of different haplotype combination (no contribution to hpd, nc-hpd region, left panel). The second type of genomic region displayed significant heterozygosity differences between isolates of different haplotype combination. In these regions, isolates with AB haplotype tended to exhibit heterozygous genotypes, while AA and BB haplotype isolates showed distinct homozygous genotypes with clear signs of recombination (medium contribution to hpd, mc-hpd region, middle panel). The third type of region was similar to the mc-hpd region but exhibited complete divergence of haplotype combinations without signs of recombination (high contribution to hpd, hc-hpd region, right panel).

**Fig.4.**
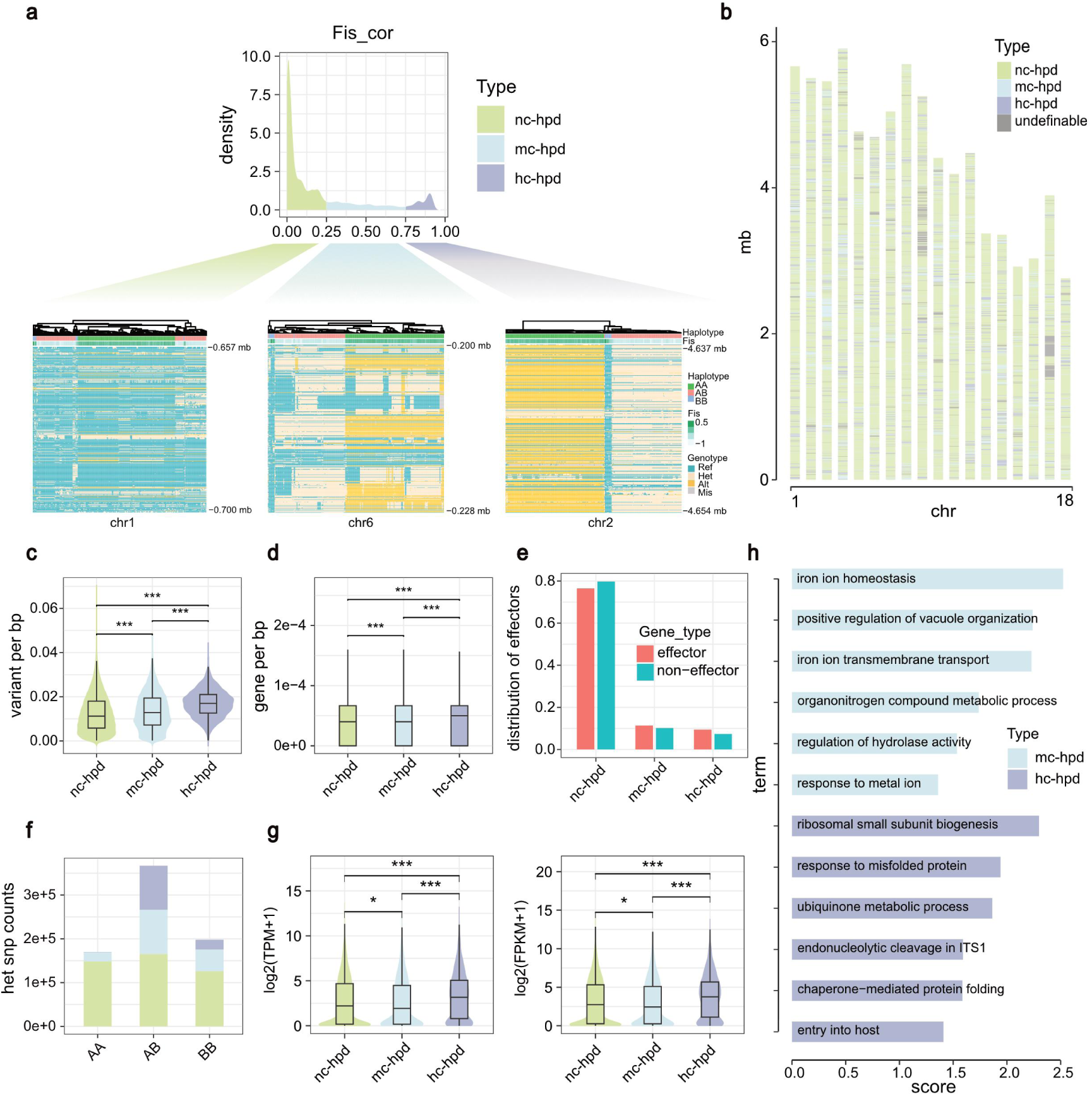
Genomic regions with three distinct landscapes of population genetic variation. a. Fis_cor was used to identify genomic regions with different population variation landscapes. The upper panel shows the distribution of Fis_cor values for all SNPs. The lower panels show genotype heatmaps for three types of regions. Each row represents a SNP, and each column represents an isolate, with rows clustered but columns ordered by chromosome coordinates. b. Distribution of the three types of genomic regions across the genome based on 5 kb sliding windows. Gray regions indicate undefinable regions. C. SNP density and d. Gene density based on 5 kb sliding windows. e. Proportions of effectors and non-effector genes. f. Numbers of heterozygous SNPs among different haplotype combinations. g. Average gene expression. h. GO term enrichment for genes located in mc-hpd and hc-hpd regions.

We created a Fis_cor index for measuring the correlation (R^2^) between SNP and heterozygosity, in order to classify different regions of the whole genome. This index could assess each SNP’s contribution to hpd. Based on the genome-wide distribution of Fis_cor, we found that setting thresholds of 0.25 and 0.75 effectively classified SNPs into different types of regions (Fig. 4a). We then performed a sliding-window statistics and split the genome into four types of chromosome regions, including 80.4% nc-hpd, 9.8% mc-hpd, 6.3% hc-hpd and 3.5% undefinable regions (too few SNPs to judge) (Supplementary Table 5). The regions of mc-hpd and hc-hpd are scattered throughout the genome (Fig. 4b), where SNP and gene are both enriched compared to nc-hpd. Furthermore, hc-hpd genomic regions tend to encode more candidate effectors compared to genomic background (Fig. 4c,d,e and Supplementary Table 6). Nc-hpd region had similar number of heterozygous SNPs among all three types of isolates, while mc-hpd and hc-hpd regions did not. AB haplotype isolates had more heterozygous SNPs in mc-hpd and hc-hpd than AA or BB haplotype isolates (Fig. 4f).

To investigate the gene expression level of these three types of regions, 250 public RNA-seq datasets from the Rust Expression Browser^30^ were aligned to the Pst134pri genome to calculate the expression of each gene. We found that genes in the hc-hpd region generally had higher expression than genes of other regions (Fig. 4g and Supplementary Table 7). Gene Ontology (GO) enrichment analysis indicated that genes in mc-hpd region were enriched in iron ion transport and signaling pathways, while genes in hc-hpd region were enriched in pathways related to protein folding, homeostasis maintenance, and host invasion. Those results indicate hc-hpd regions are crucial for the virulence of the pathogen (Fig. 4h and Supplementary Table 8).

In summary, these three types of regions showed distinct population variation landscapes, with mc-hpd and hc-hpd regions contributing to A and B haplotype divergence and hpd. Notably, the hc-hpd is similar to introgression segments, suggesting its potential role as homologous segments introgressed into the genome of *Pst*. Looking back at the previous puzzling results in the first section of this paper, we can now answer them. The bimodal distribution of minor allele frequency (MAF) is due to the different distributions of MAF in three types of regions. The MAF of mc-hpd and hc-hpd concentrated between 0.2 and 0.3 (Supplementary Fig. 4a). The peak of F_MISS at 0.36 is caused by the absence of B haplotype-specific segments in AA haplotype isolates. Interestingly, many of these haplotype-specific segments belong to mc-hpd region (Supplementary Fig. 4b,c). Similarly, AB haplotype isolates have fewer missing sites, while AA and BB haplotype isolates have more missing sites (Supplementary Fig. 4d). Understanding the PCA and phylogenetic tree clustering results through haplotype combinations seems more logical (Supplementary Fig. 4e,f). The estimation of ancestral components further supported this, when K=2, the result showed that AA and BB haplotype isolates have different single ancestral components, however AB haplotye isolates represent a mixture of the two (Supplementary Fig. 4g).

### A/B haploid-inherited regions

As shown in Fig. 4a, mc-hpd genomic regions shows prominent pattern of recombination, while hc-hpd does not. In order to understand the relationship between the A-B divergence and sexual reproduction, we calculated and compared linkage disequilibrium (LD) decay within and between species. *Magnaporthe oryzae* (synonym of *Pyricularia oryzae*), a soil-borne pathogen which is thought to be lack of sexual reproduction in nature^39^. It could be used as a negative control for pattern observed in genotype heatmap of *Pst*, which has confirmed sexual reproduction in the natural environment. We collected 183 WGS public data of *M. oryzae*^39-46^ and performed SNP calling using Pop131 as the reference genome (Supplementary Table 9). LD decay analysis revealed that *Pst* populations exhibited faster decay and lower baseline compared to *M. oryzae*, indicating more frequent recombination events in *Pst* (Supplementary Fig. 5a). Differences in LD decay were also observed between subpopulations of different haplotype combinations. LD decay in subpopulation of AA haplotype showed a lower baseline compared to AB haplotype, suggesting higher recombination rates in AA haplotype (Fig. 5a).

**Fig.5.**
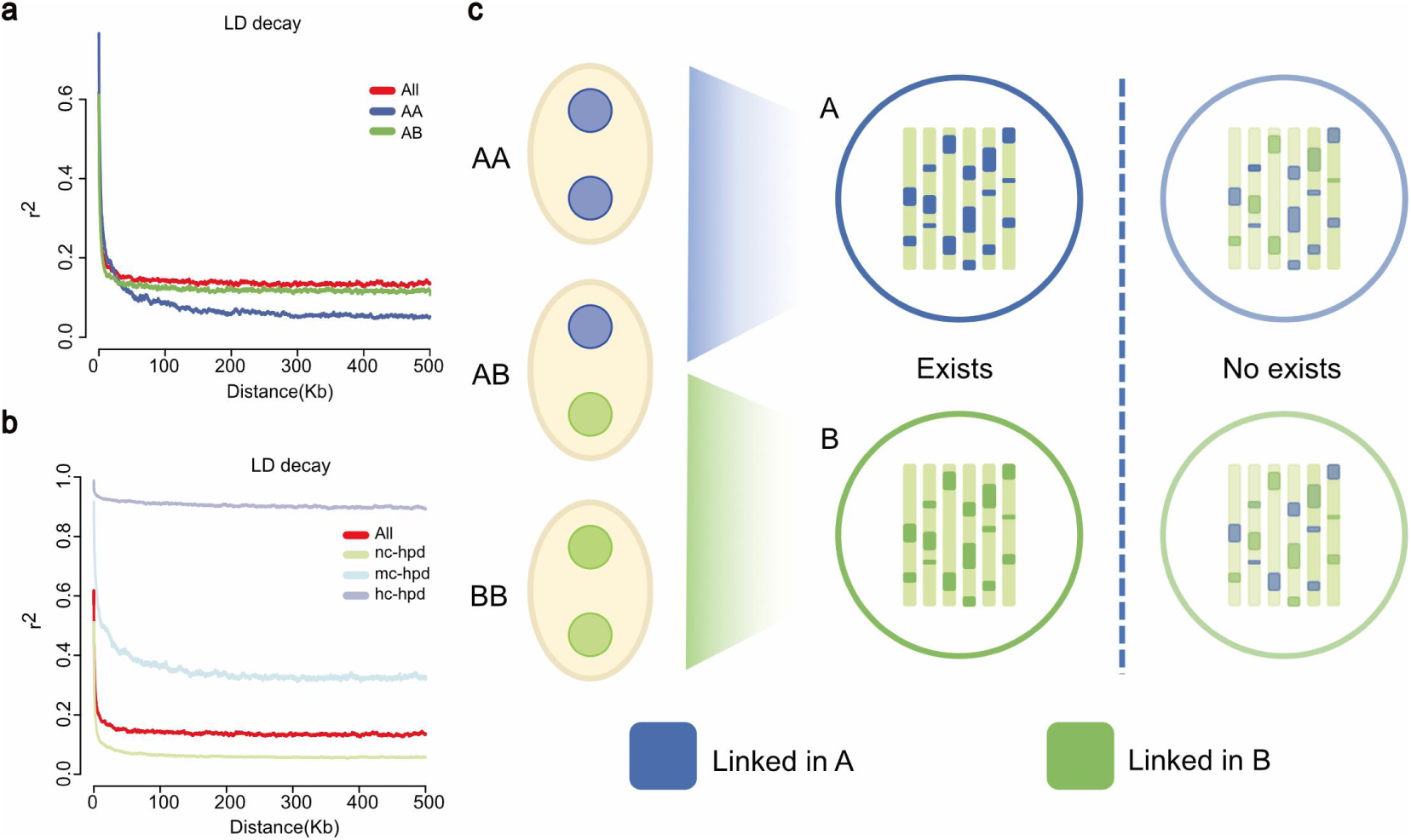
A/B haploid-inherited regions with rare recombination. a. LD decay was performed for subpopulation defined by haplotype combinations, with different colors representing distinct haplotype combinations (AA, AB). b. LD decay was performed for the three types of regions. In hc-hpd regions, LD does not decay significantly, indicating long-distance linkage. c. A schematic of the A/B haploid-inherited region: these haplotypes exhibit inter-chromosomal linkage in A/B haploid. Genomic segments inherited from the A haplotype and from the B haplotype have not recombined.

The Asian subpopulation showed the fastest decay, followed by the North American subpopulation, and the European subpopulation showed the slowest decay with progressively higher LD baselines. This is consistent with the different geographical distribution of haplotype combinations, such as AA haplotype isolates being more prevalent in Asia (Supplementary Fig. 5b). LD decay also differed across genomic regions. The most notable observation is that the LD in hc-hpd regions does not decay with increasing chromosome physical distance. It maintains a baseline of around 0.9, indicating long-range linkage (Fig. 5b).

Moreover, we confirmed that hc-hpd regions exhibit inter-chromosomal linkage, suggesting that these regions are inherited by haploids. A and B haploids consistently maintain their respective haplotype in the hc-hpd regions without recombination between themselves (Supplementary Fig. 5c, Fig. 5c). This may suggest that genes within these regions have established fixed epistatic networks that regulate essential biological functions, therefore the interchange of gene nodes between A and B networks could not be allowed. Furthermore, in the AB haplotype subpopulation, the A/B haploid-inherited region consistently remains heterozygous and does not undergo recombination, which increases the LD decay baseline.

### Sexual reproduction promotes the evolution of virulence in *Pst*

The contrasting pattern in terms of recombination for hc-hpd and mc-hpd highlighted the effectiveness of our data analyzing strategy in capturing genomic evidence for alternative life cycle. Sexual reproduction events of *Pst* in nature are challenging to capture. However, prior studies successfully induced sexual reproduction of *Pst* in controlled environments. Notably, Chen et al. (2020) created an inbred population containing 117 progeny isolates^29^ and published phenotype and WGS data, initially used for mapping avirulence (*Avr*) genes. We aim to re-utilize these data to understand the genomic recombination patterns during sexual recombination in *Pst* and their association with virulence evolution.

First, we removed 25 isolates with higher heterozygosity compared to the parent (Supplementary Fig. 6a). Using Pst134pri as the reference genome, we employed Dfseq-calling to generate a genetic variation map for the inbred population (Supplementary Table 10). Upon examining the phenotypes of the inbred population, we noted virulence segregation among the progeny isolates. The parental isolates exhibited weak virulence against *Yr7*, *Yr8*, *Yr43*, *Yr4*4, and *YrExp2* single-gene lines, whereas progenies with stronger virulence were identified. Through GWAS re-mapping, we mapped signals for *AvrYr43*, *AvrYr44*, *AvrYr7*, and *AvrYrExp2* to the intervals chr3: 3.10-3.11 Mb and chr7: 0.04-0.14 Mb. Signals for *AvrYr8* were detected near chr9: 5.02 Mb and chr18: 0.27 Mb (Fig. 6a and Supplementary Fig. 6b and Supplementary Table 11).

**Fig.6.**
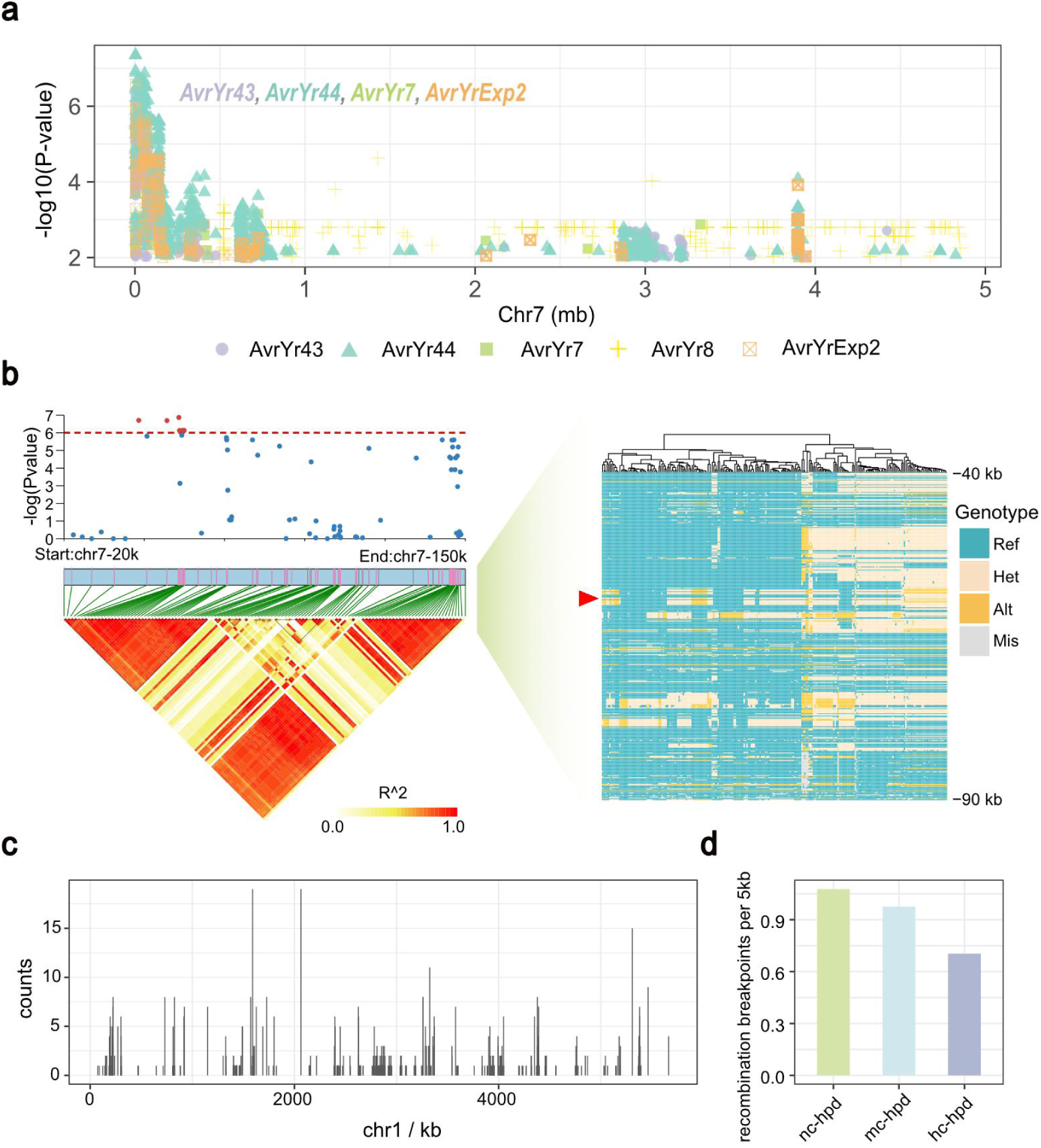
Avr gene re-mapping and recombination breakpoint. a. GWAS re-mapping results revealed a cluster of *Avr* genes on chromosome 7, including signals for *AvrYr7*, *AvrYr43*, *AvrYr44*, and *AvrYrExp2*. b. The left panel shows the LD block for the candidate interval on chromosome 7 in the inbred population. The right panel displays the genotype of this candidate interval in natural populations. Each row represents a SNP and each column represents an isolate, with rows clustered but columns ordered by chromosome coordinates. Clear recombination fingerprints in this interval among natural population indicate that sexual reproduction driven the evolution and diversity of *Pst* virulence. c. Distribution of recombination breakpoints along chromosome 1 in all progenies, with the x-axis showing physical position and the y-axis showing the frequency of breakpoints at each location. d. Density of recombination breakpoints in three types of regions. In hc-hpd regions, fewer recombination breakpoints are observed, suggesting a lower frequency of recombination in these regions.

Large haplotype blocks were discovered in the candidate interval chr7: 0.02-0.15 Mb (Fig. 6b). The genotype heatmap revealed various haplotypes with clear recombination fingerprints for this interval in our natural population panel. By checking the relationship between recombination and virulence changes in this inbred population, we found that the transition of virulence loci from heterozygous to homozygous could help *Pst* enhance its virulence. It indicated that sexual reproduction in the natural environment may facilitate *Pst*’s virulence evolution, thus better overcoming host resistance (Supplementary Fig. 6b and Supplementary Table 12).

Subsequently, we analyzed the distribution of recombination breakpoints of all progeny isolates (Supplementary Table 13) and found a notable preference. More breakpoints cluster in nc-hpd region and fewer in hc-hpd region (Fig. 6c,d and Supplementary Fig. 7). This partially supports the hypothesis that hc-hpd serves as an A/B haploid-inherited region subject to recombination suppression.

### A global *Pst* genotype display platform

For a long time, researchers sought for efficient methods to capture the global migration and diversity of pathogen^26,47^. It is essential for predicting disease outbreaks and epidemics, allowing for the timely disease management. Here, we built a user-friendly platform for visualizing global *Pst* genotype heatmap. This platform hosts the genetic variation map of 239 global *Pst* isolates gathered in this study (Supplementary Fig. 8). Accessing the platform, users can select specific isolates and genomic regions, enabling precise retrieval of any genotype information in the dataset. The platform also allows users to download genotype heatmaps generated through it. On the one hand, we are confident that this genotype information is invaluable for understanding the findings of our study. We aim to develop this platform to better present our discoveries and invite readers to verify the credibility of our research through it. On the other hand, we believe this platform can assist researchers to better understand the diversity and evolutionary dynamics of *Pst.* It provides a solid data foundation for studying the virulence evolution of crop fungi pathogen.

## Discussion

For a long time, pathologists achieved significant results through conducting extensive researches on issues such as strain migration, virulence evolution, and population diversity^37,48,49^. From traditional molecular markers to high-throughput sequencing data^50^, advances in sequencing technologies provided new opportunities for these studies, allowing researchers to deepen their understanding of pathogens. However, due to limitations in sampling and high strain cultivation costs, population studies of pathogens remained scarce. Due to lack of quality control, errors like strain mixing also hindered research progress. There is an urgent need to develop new platforms and methods to facilitate research development. New methods should be suitable for microbial population studies. They could be rapid sampling and strain identification methods, low-cost strain purification and preservation techniques, and computational methods for detecting strain mixing or contamination. In recent years, researchers created some innovative methods. Hubbard et al. established field pathogenomics, which eliminated the laboratory strain purification process by directly sequencing infected leaves in the field, significantly improving sampling efficiency^26^. Radhakrishnan et al. launched the Marple platform, a point-of-care, strain-level disease diagnostics and surveillance tool that can quickly capture strain migration. It costs a very low price and is easy to deploy^47^. Our study also introduces a new dikaryotic microorganisms variant detection pipeline, Dfseq-calling. By using Dfseq-calling, we can easily detect potential strain mixing events in isolate sample through short reads data. After filtration, it could provide high-quality genetic variation information, ensuring accurate downstream analysis and facilitating the study of population genetics in *Pst*. As sequencing cost continues to decrease, more isolates will be sequenced by the community. Those data will be available through public repositories such as NCBI SRA. Our work here showed the power of large-scale public data mining in revealing the evolutionary dynamics of crop pathogen and provided an example pipeline for future studies.

Sex is considered to influence the evolution of species^51^. Many microorganisms exhibit multiple reproduction strategies. For example, rust fungi are thought to reproduce through asexual, sexual, and parasexual cycles^1^. However, the evolutionary significance of such alternative reproduction strategies in rust fungi remains unclear. Some researchers suggest that the hpd in *Pst* is caused by differences in reproductive strategies across strains. As the Meselson effect describes, high-heterozygosity strains are believed to be generated through long-term asexual reproduction process, whereas frequent sexual reproduction produces low-heterozygosity strains^7-20^. At population level, our findings prove that hpd arises from different haplotype combinations. Specifically, it is driven by suspected introgression segments, which have not undergone extensive recombination. These segments remain heterozygous in AB haplotype isolates, leading to heterozygosity differences between AA/BB and AB haplotype isolates. This conclusion is similar to experiments on bdelloid rotifers^21,22^, where heterozygosity was linked to the introgression of paralogous genes. However, we did not identify a clear donor for such introgression segments. This hypothesis is still challenged. Nevertheless, we still want to emphasize the unreliability of inferring a strain’s reproductive history solely based on estimates of genomic heterozygosity. We observed clear fingerprints of sexual recombination even in highly heterozygous isolates. Previous studies suggested that heterozygosity can be introduced not only through asexual reproduction but also via hybridization^52^. Therefore, we argue the judgments of regarding an isolate’s reproductive strategy should be based on field observations and experiments.

Schwessinger et al. studied a low-heterozygosity strain DK-0911, which had been outbreak in Europe in recent years^12^. This strain belongs to the PstS7 lineage (warriors), which spread widely across Europe after 2010 and is considered more similar to strains from the Himalayan region that undergo frequent sexual reproduction. It is thought to be long-term inbreeding. This study discussed the heterozygosity levels of several assembled strains and the impact of different reproduction strategies on genome adaptability. It mentioned that DK-0911’s genome contains gene pairs (alleles in the two haploid nuclei) with different variation landscapes and evolutionary rates. It resembles the three types of regions with different population variation landscapes we observed in *Pst*, suggesting introgression of intra- and inter-species segments. Schwessinger et al. proposed an explanation for the different evolutionary rates. The first situation involves a steady accumulation of mutations during asexual reproduction, leading to a gradual and linear increase in differences between gene pairs. The second situation involves variation introduced at a specific moment, such as through mating between genetically distinct individuals or somatic recombination via the transfer of a single nucleus. Our study supports the second situation at the population level, suggesting that gene pairs with faster evolutionary rates and higher heterozygosity are due to the introgression of intra- and inter-species segments. Notably, our genetic variation map also includes DK-0911, which we classified as an AA haplotype, significantly different from the AB haplotype isolates found in Europe. Our research supported some aspects of previous hypotheses at the population level while highlighting the one-sidedness of other conclusions, advancing the understanding of dikaryotic pathogen evolution.

Our study defines the worldwide-prevalent haplotypes A and B. We found that certain regions (hc-hpd regions) exhibit differentiation between A and B haplotype while maintaining haploid-inherited and displaying low recombination levels. These regions are responsible for sustaining the divergence of A and B haplotypes. However, we have no clues regarding how this differentiation occurs. We can only confirm that the distribution of these regions is scattered across the genome and similar to introgression segments. The variation landscape does not resemble linear haploid differentiation resulting from long-term asexual reproduction in a monophyletic lineage. A rational hypothesis is that the B haplotype (or A haplotype) is a product of hybridization between the A haplotype (or B haplotype) and C haplotype. This hybridization event incorporated exogenous segments from the C haplotype into the background of the A haplotype (or B haplotype). Previous studies induced hybridization events of different rust fungi in experimental environment^53^, lending some reliability to this hypothesis. C haplotype isolates may be stripe rust fungi infecting different hosts or other species of rust fungi. We illustrated this hypothesis, proposing that the B haplotype is a product of hybridization between A haplotype and C haplotype. Then, it leads to the formation of AB haplotype isolates, which possess greater adaptability and spread globally (Supplementary Fig. 9). The validity of this hypothesis requires further verification through more sampling and sequencing of isolates, including additional BB haplotype isolates and more different forma specialis isolates, particularly isolates hosted on Poaceae. Although the accuracy of this hypothesis is open to question, the divergence of A and B haplotypes is undoubted. This observation prompts us to focus on inter- and intra-species hybridization events that occur in natural environment among rust fungi, including sexual reproduction and somatic hybridization. These events may be crucial ways for pathogens to acquire adaptability and novel virulence^54^.

Our study identified A/B haploid-inherited regions, which are scattered across the genome. They are inter-chromosomal linkage, maintaining differentiation between A and B haplotypes without recombination. The genes located in these regions exhibit higher expression levels, suggesting their importance in supporting essential life processes of *Pst*. The existence of such regions reminds us to focus on the haploid stages of *Pst* (basidiospores and aeciospores). On the one hand, there are compatibility issues between haplotypes during mating. On the other hand, it is still unknown how the selection pressure imposed by the host on the haploid generation affects the *Pst* genome. These A/B haploid-inherited regions might be selected during the haploid stage. Maintaining their integrity could ensure each haploid isolate survives independently. Although this genomic landscape is intriguing, it poses challenges for *Avr* gene mapping. Genes in A/B haploid-inherited regions are linked across chromosomes. If *Avr* genes are located within these region, GWAS would be difficult to identify them. Similar to how population stratification affects GWAS, non-*Avr* genes within A/B haploid-inherited regions may create false positive signals. We still lack clues on why these regions maintain A/B haploid-inherited or the validity of the concept itself. A promising approach to address this would be creating an inbred population from an AB haplotype isolate parent or a hybrid line between AA and BB haplotype isolates. Then sequencing their progeny and conducting further experiments. Exploring the biological mechanisms behind it might help us understand how *Pst* maintains virulence and ensures survival.

In conclusion, our research challenged traditional views on the causes of high heterozygosity in *Pst*. It highlighted the interesting event of A and B haplotype divergence and its contribution to the global diversity and environmental adaptability in *Pst*. We accurately answered the causes of hpd and provided insights into how alternative reproduction strategies influence *Pst* evolution. While some hypotheses presented here may be speculative and subject to scrutiny, they offer valuable clues for future studies on the origin, evolution, and virulence evolution of *Pst*.

## Method

### Data source and quality control

The genome assemblies Pst134 and AZ2 were acquired from NCBI under BioProject numbers PRJNA1026770 and PRJNA749614. The protein sequences (FASTA format) and gene annotations (GFF format) were gathered for each assembly. Worldwide *Pst* WGS data^12,25-33,35-37,55^ were under 18 NCBI BioProjects, with additional data from the China National Genomics Data Center under BioProject number PRJCA014799. Detailed accession numbers and BioProject numbers are listed in Supplementary Table 1. *Pst* RNA-seq data^30^ were acquired entirely from NCBI under BioProject numbers PRJEB39201 and PRJEB31334, with details in Supplementary Table 7. The WGS data for *M. oryzae*^39-46^ were under 11 NCBI BioProjects, with accession numbers and BioProject numbers listed in Supplementary Table 9. Prior to alignment, all WGS data were quality-controlled using fastp^56^ with default parameters.

### Dfseq-calling pipeline

The Dfseq-calling pipeline is an improved version of the SNP calling workflow. It was designed to address mistakes posed by low-quality, dikaryotic microbial sequencing data through additional filtering steps. The detailed steps include:

1. Initial Filtering: Raw data is filtered using fastp with default parameters.
2. Alignment: Reads are aligned to the haploid reference genome using BWA-MEM^57^ with default parameters, generating SAM files.
3. Conversion and Deduplication: SAM files are converted to BAM format and deduplicated using Samtools^58^.
4. Calling: SNPs are called using FreeBayes 1.3.5^59^ with the parameters-use-best-n-alleles 6-ploidy 2 in parallel mode, producing VCF files.
5. TE Region Masking: Transposable elements (TEs) are identified using RepeatMasker (http://www.repeatmasker.org). Mask and filter out SNPs within TE regions.
6. Strain-Mixing Isolates Filtering: SNPs’ DP (depth) and RO (number of reference observations) values are extracted from the VCF to calculate AD (allele balance, AD = RO/DP). Isolates with heterozygous SNPs that deviate significantly from the normal distribution (μ=0.5) are flagged as potentially strain-mixing isolates, being filtered.
7. Regular Filtering: The VCF is further filtered with bcftools^60^, using bcftools -i ’QUAL > 20 & SAF > 0 & SAR > 0 & RPR > 1 & RPL > 1 & AC > 0’ to retain high-quality SNPs, excluding multi-allelic sites.

Using this pipeline, two genetic variation maps were generated: a map based on 239 worldwide *Pst* isolates and another for inbred population. For the *M. oryzae* genetic variation map, a simpler SNP calling pipeline was employed, omitting steps 5 and 6.

### Genetic diversity analysis

The MAF was calculated using vcftools^61^ --freq. The F_MISS was calculated using vcftools --miss-site. The Fis was calculated using vcftools --het. IBS between isolates was assessed using vcftools --relatedness. Fst was calculated using vcftools --weir-fst-pop. To construct an accurate phylogenetic tree of *Pst*, we selected SNPs from gene regions for tree building. The phylogenetic tree was built using iqtree^62^ with parameters -m GTR+F+G4+ASC -fast -nt 50 -st DNA. Ancestral component was estimated using admixture^63^ with default parameters. PCA was performed on standardized genotype data using R’s prcomp function with default parameters. We selected SNPs from gene regions for LD calculation. LD was calculated using PopLDdecay-3.42^64^, with parameters -maxdist 500, and plots were generated using Plot_MultiPop.pl. MAF, F_MISS, Fis, IBS, and Fst distributions were visualized using ggplot2^65^. The phylogenetic tree was visualized with iTOL^66^, and the ancestral component was plotted using pophelper^67^.

### Genome comparisons and k-mer screening

The four haploid genomes were compared to each other with MUMmer-4.0.0rc1^68^, using nucmer. After alignment, we applied the filtering with mummer-4.0.0rc1 delta-filter using parameters -1 -q -r and used show-snps -Clr to count SNPs between four haploid genomes. The collinearity dot plot was generated using D-GENIES^69^, with Minimap2 v2.28 as the aligner and the parameter -repeatedness set to “some repeats.” Mash was used for k-mer containment screening. We screened the four haploid genomes with sketch settings -s 500000 -k 32, returning the distance and shared k-mers counts between genomes and sequencing reads. The shared k-mer counts distribution was visualized with ggplot2 in R, and the heatmap of shared k-mers was drawn using ComplexHeatmap^70^.

### Classification and analysis of genomic regions with different landscape of population variation

We defined the Fis_cor index to classify different types of SNPs, further enabling the definition of chromosome regions. Fis_cor evaluates each SNP’s contribution to hpd. First, we assigned values to each SNP: 1 for heterozygous and 0 for homozygous. Next, we calculated the Pearson correlation coefficient (R^2^) between each SNP’s value and each isolates’ Fis using the cor function in R, similar to the concept of GWAS where Fis is treated as trait. SNPs with 0.25 >= R^2^ > 0 were classified as variants that do not contribute to hpd (nc-hpd), those with 0.75 >= R^2^ > 0.25 as medium contribution (mc-hpd), and R^2^ > 0.75 as high contributors (hc-hpd). We applied a sliding window with a 5 kb size, moving 1 kb each time, to detect the distribution of different SNP types. If a type of SNP is more abundant within a window, the window region is defined as that SNP type. Otherwise, for windows with no SNP, it would be categorized as an undefinable type region. SNP and gene density were calculated within these windows. We predicted effectors using SignalP 5.0 and EffectorP 3.0^71,72^. For gene expression analysis, we aligned transcriptome reads to the Pst134pri genome using STAR^73^ and computed TPM and FPKM values via featureCounts^74^. For GO enrichment analysis of different regions, we annotated the entire gene set with EggNOG v2^75^, selecting fungi for the taxonomic scope, and performed enrichment analysis using TBtools^76^. We used ggplot and pheatmap for visualization.

### Re-mapping of sequencing data from a published inbred population

We gathered 119 inbred population isolates WGS data^29^ from NCBI under BioProject number PRJNA599033. We performed snp calling using Dfseq-calling pipeline, removing isolates with abnormal heterozygosity or potential strain-mixing risk. The VCF files were converted into hapmap format. Using phenotypic data provided by Chen et al.’s research, we conducted a GWAS (Genome-wide association study) using GAPIT with parameter pca.total set to 3. After mapping, we visualized the haplotype blocks of these candidate intervals with LDBlockShow^77^. The Manhattan plots were generated using ggplot, and genotype heatmaps of the candidate interval were drawn using pheatmap.

### Finding recombination breakpoints

We wrote an R script to calculate the recombination breakpoints for each progeny (identify the boundary between homozygous and heterozygous genotype regions as a recombination breakpoint). We calculated the density of recombination breakpoints in three types of regions (per 5 kb) and visualized the distribution using ggplot.

### Accessibility of genetic variation map data

After constructing a worldwide *Pst* genetic variation map, we made it accessible to readers through multiple platforms (Supplementary Fig. 8). The data was uploaded to the European Variation Archive (EVA) and Genome Variation Map (GVM)^78^. Additionally, we built a genotype display platform using Shiny and PostgreSQL, which can be accessed at http://crop-pathogen-genomics.cn/pst_gt_heatmap/.

## Funding

This work was supported by the Biological Breeding-Major Projects (2023ZD04076), the National Key Research and Development Program of China (2023YFF1000100) and the Research Program for Network Security and Information of the Chinese Academy of Sciences (CAS-WX2021SF-0109).

---

**Fig.S1.**
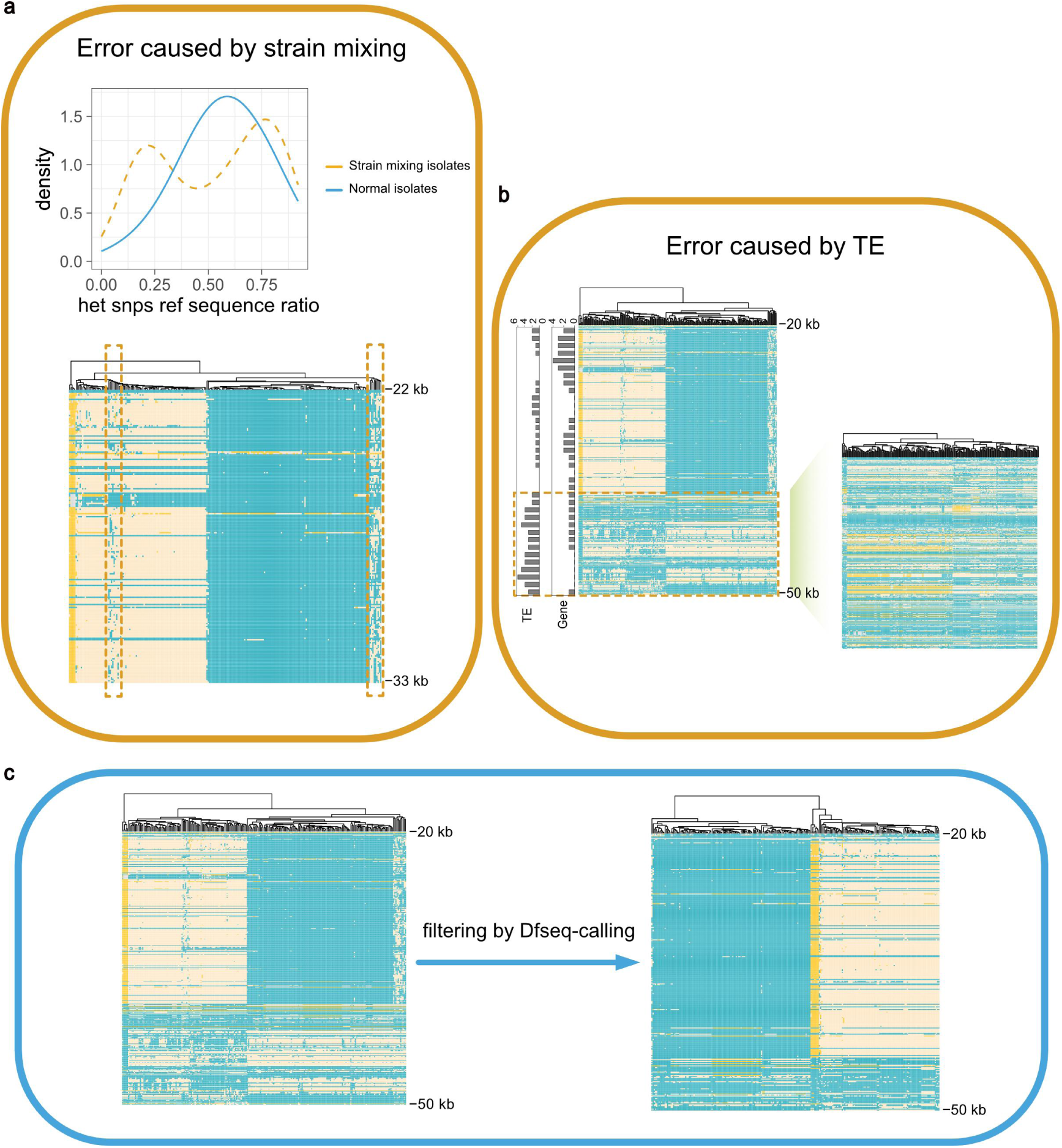
Errors during traditional SNP calling process and the cleaner genetic variation map obtained after filtering by Dfsep-calling pipeline. Taking the region from chr1 20-50kb as an example. Each row represents a SNP and each column represents an isolate, with rows clustered but columns unclustered. a. SNP calling errors caused by strain mixing risk. By calculating the AD (allele balance) values of heterozygous SNPs in each strain, we identified some isolates with an abnormal bimodal distribution of AD values, indicating strain-mixing. This error is visually evident on the genotype heatmap. The dashed-line-marked isolates exhibit a mosaic genotype, rendering the data highly unreliable. b. SNP calling errors caused by TEs. The left panel highlights a region marked with dashed lines, showing a similar mosaic genotype of linkage loss. Further analysis revealed that these regions are enriched for TEs but gene-poor. The right panel presents a genotype heatmap of all SNPs in TE regions on chromosome 1, displaying a widespread mosaic genotype pattern, indicating low data reliability in these regions. c. Using chr1 20-50kb as an example, the genotype shows a cleaner linkage pattern after filtering by Dfseq-calling pipeline.

**Fig.S2.**
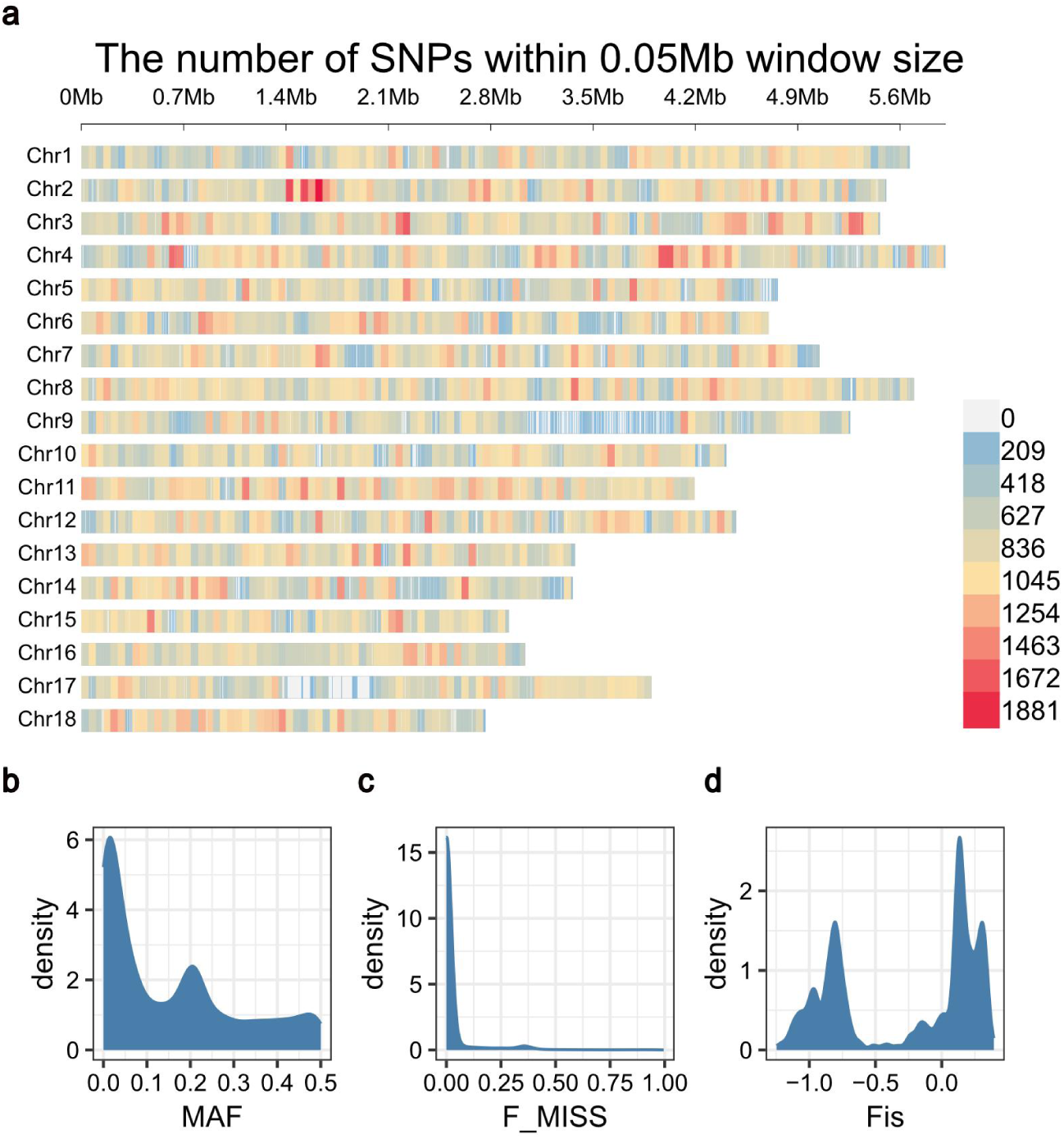
Population statistics of the global *Pst* genetic variation map. a. The SNP density distribution of the global *Pst* genetic variation map. b. The minor allele frequency (MAF) distribution of SNPs in the global *Pst* genetic variation map. c. The F_MISS distribution of SNPs in the global *Pst* genetic variation map. d. The Fis distribution of global *Pst* isolates.

**Fig.S3.**
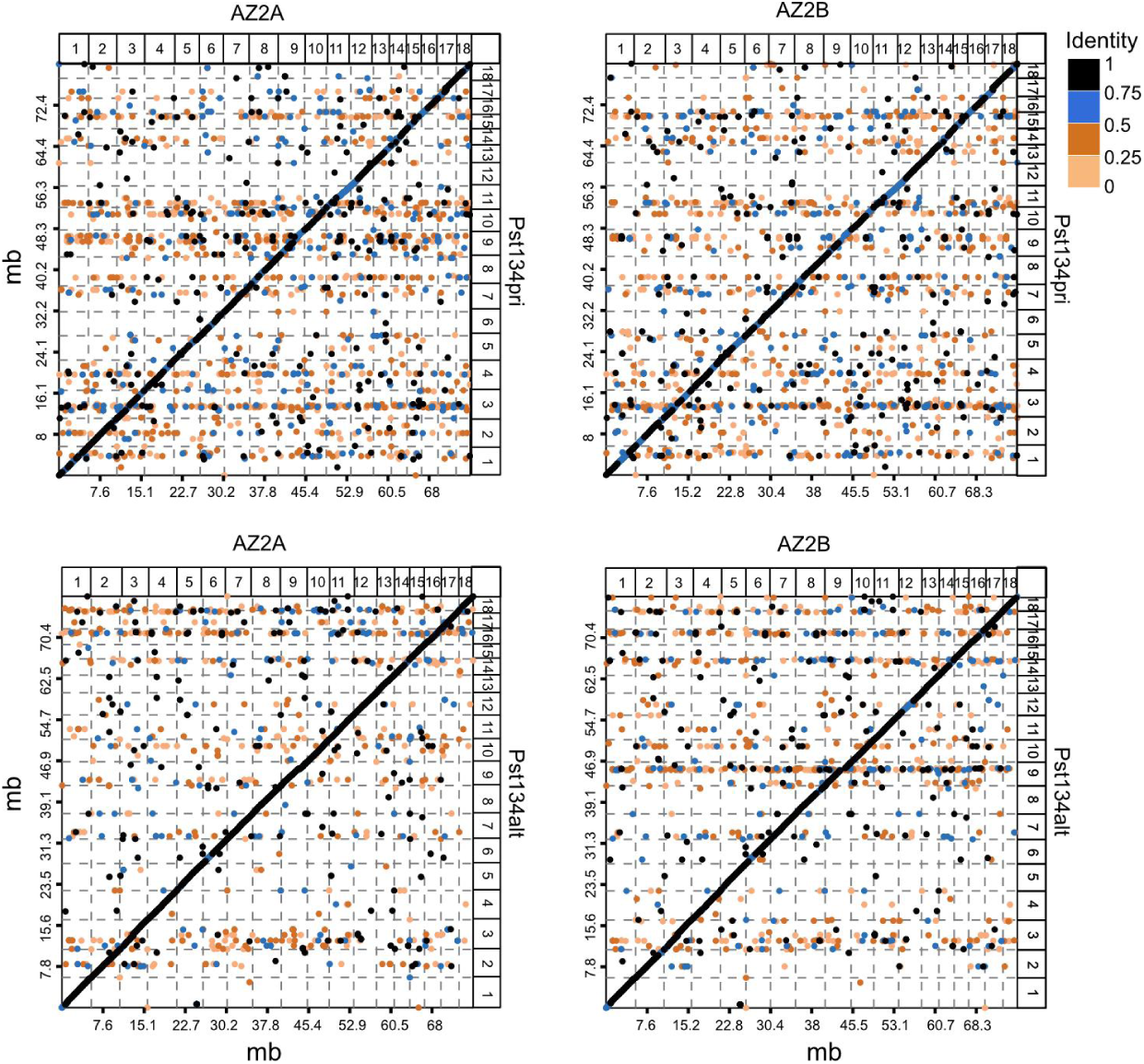
Comparison of multiple haploid genomes of Pst: AZ2A-Pst134pri, AZ2B-Pst134pri, AZ2A-Pst134alt, and AZ2B-Pst134alt. Point and line colors indicate sequence identity.

**Fig.S4.**
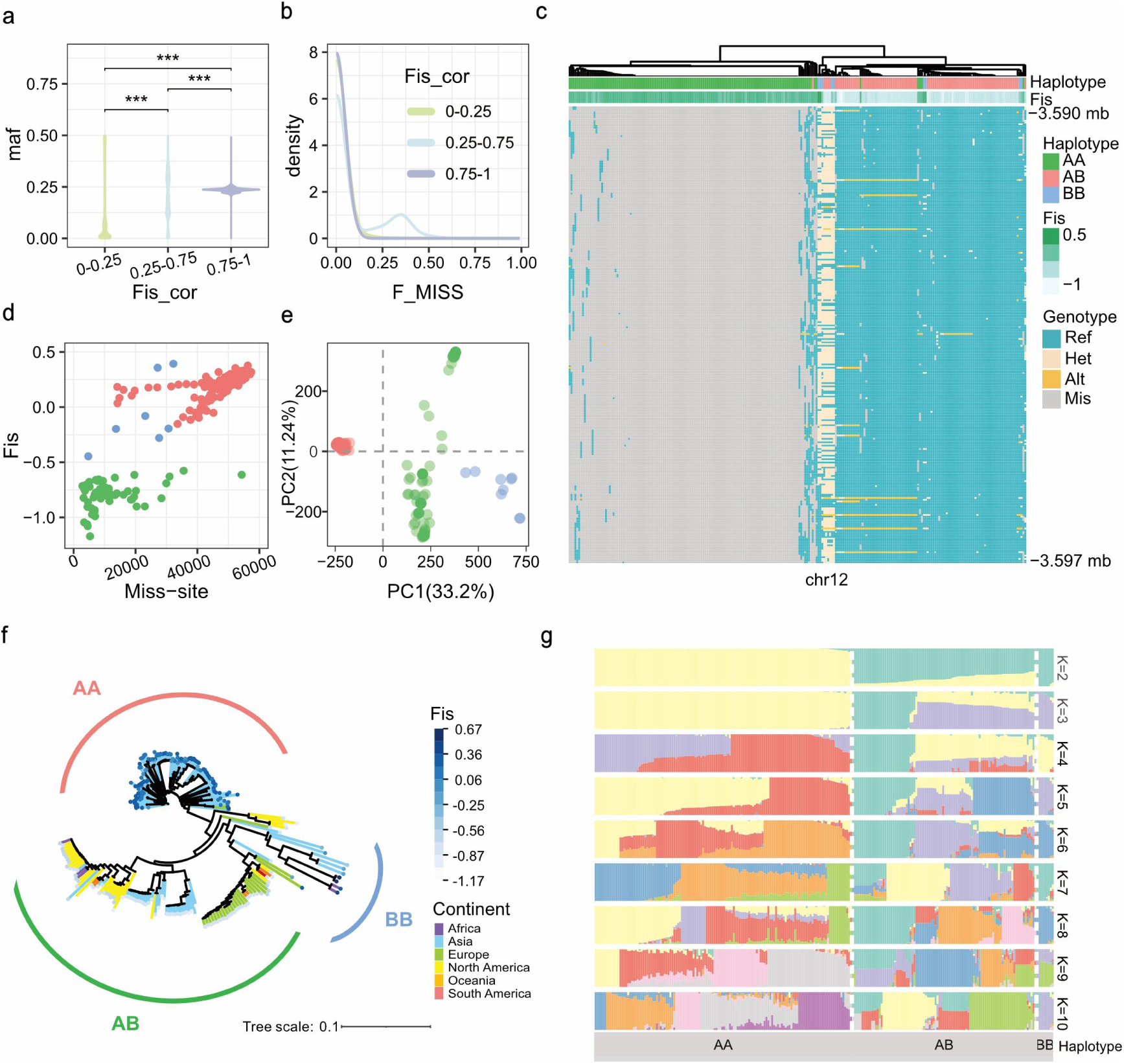
Using A/B haplotype hypothesis to explain the population variation and genomic characteristics of *Pst*. a. The MAF distribution of SNPs in three types of regions: nc-hpd, mc-hpd, and hc-hpd regions. b. The F_MISS distribution of SNPs in the three regions. c. Using the 3.590-3.597 Mb region on chromosome 12 as an example, a segment that is present in the B haploid genome but absent in the A haploid genome. This segment is missing in AA haplotype combination isolates but is present in AB and BB haplotype combination isolates. d. The relationship between the number of missing sites and heterozygosity. AA haplotype isolates have more missing sites, while AB haplotype isolates have fewer. e. PCA of global *Pst* isolates viewed from the perspective of haplotype combinations. The colors represent haplotype combination types. The same haplotype combination isolates clustered together. Results also showed significant divergence between AA, AB, and BB haplotype combination isolates. f. The phylogenetic tree of global *Pst* isolates viewed from the perspective of haplotype combinations, where AA, AB, and BB grouped distinctly. g. Ancestral component analysis of global *Pst* isolates viewed from the perspective of haplotype combinations. When K=2, AA and BB isolates exhibit distinct single ancestral components, while AB isolates show a mix of the AA and BB ancestral components.

**Fig. S5.**
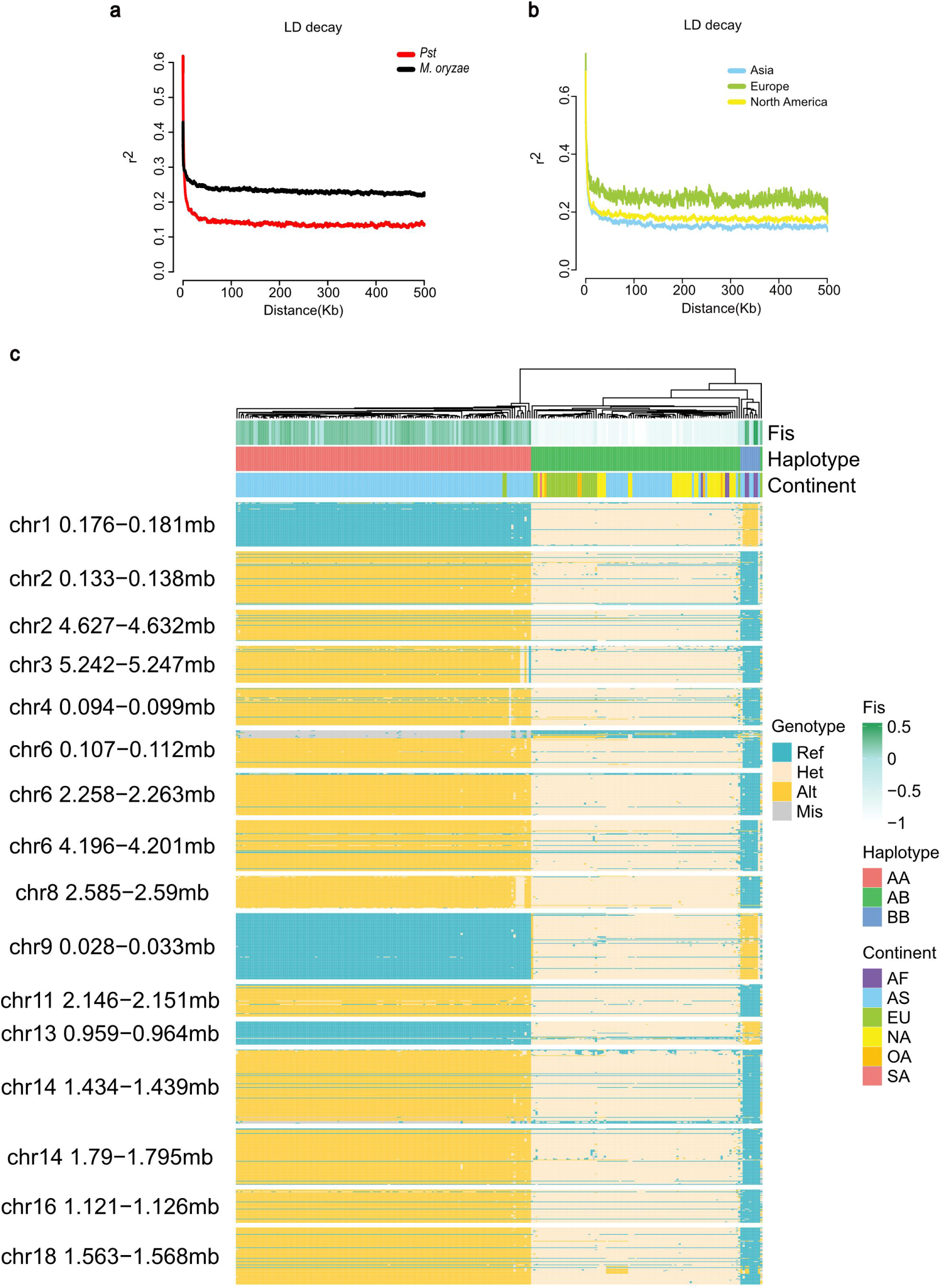
LD decay and the genotypes of A/B haploid-inherited regions. a. A comparison of LD decay between *Pst* and *M. oryzae*. b. A comparison of LD decay among *Pst* isolates from different continents. c. Genotype heatmap of several A and B genome segments exhibiting inter-chromosomal linkage. Physical positions labeled on the left. Each row represents a SNP and each column represents an isolate, with rows clustered but columns ordered by chromosome coordinates.

**Fig. S6.**
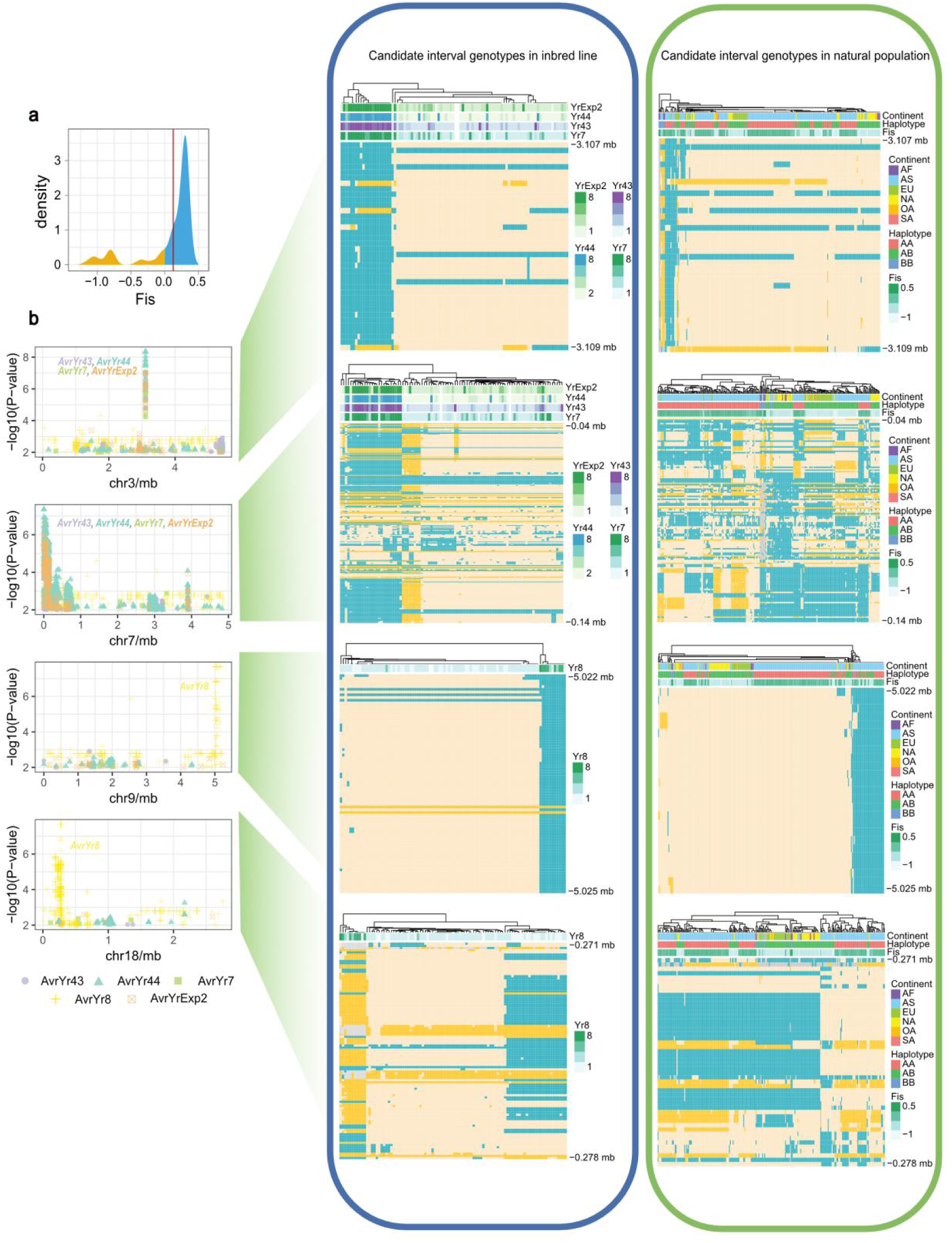
Candidate intervals of *Avr* candidate genes. a. The distribution of Fis in inbred population isolates. The red line indicates the Fis of parental isolate. Some progenies exhibit increased heterozygosity after inbreeding, and these isolates were removed. b. Genotype heatmap for four candidate intervals in both the inbred and natural populations. The Manhattan plot on the left indicates these four candidate regions. The genotype heatmap on the right includes both the inbred and natural populations. The top annotation provides information about isolates. Each row represents a SNP and each column represents an isolate, with rows clustered but columns ordered by chromosome coordinates.

**Fig.S7.**
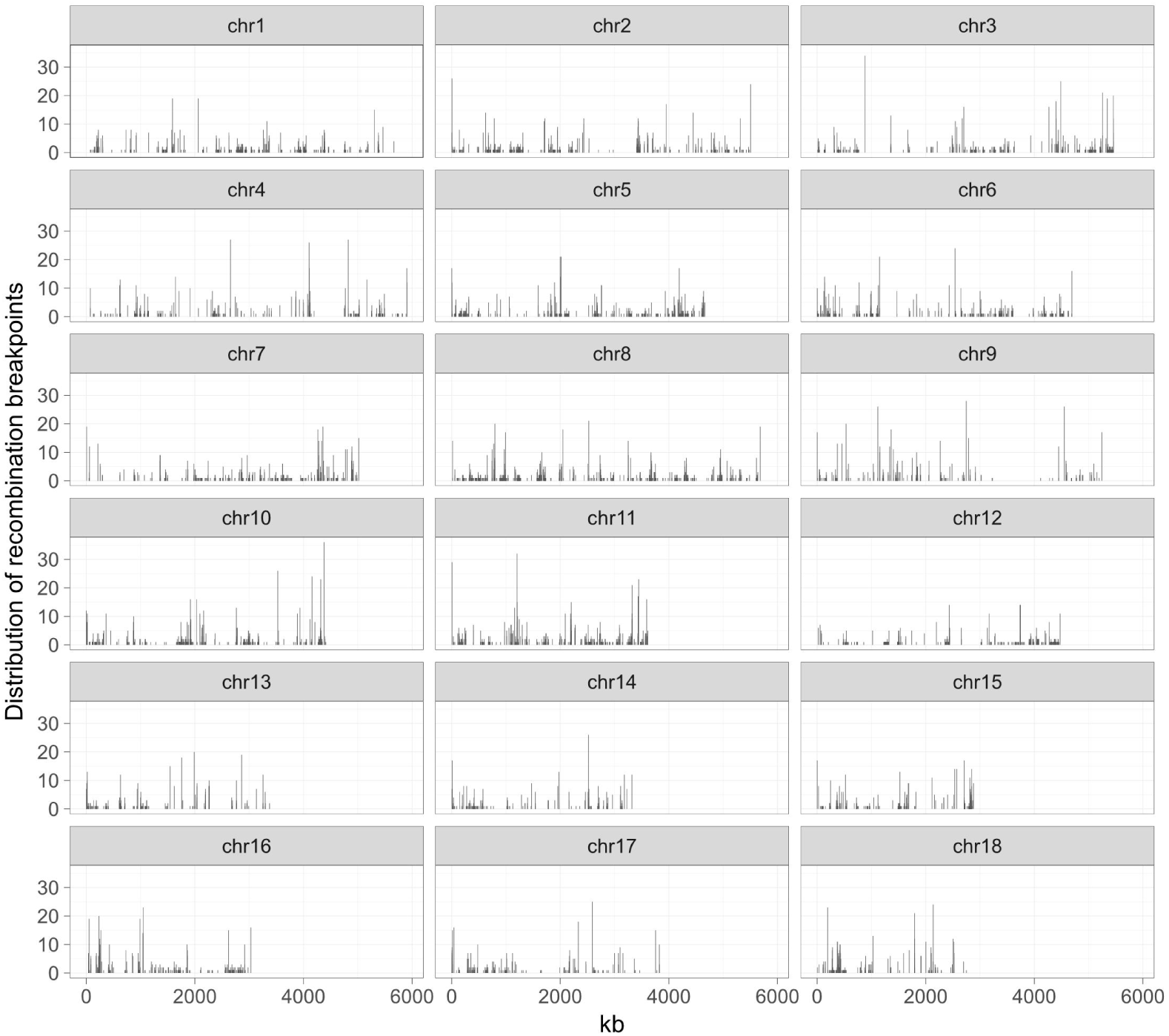
The distribution of recombination breakpoints in the progeny of the inbred population.

**Fig.S8.**
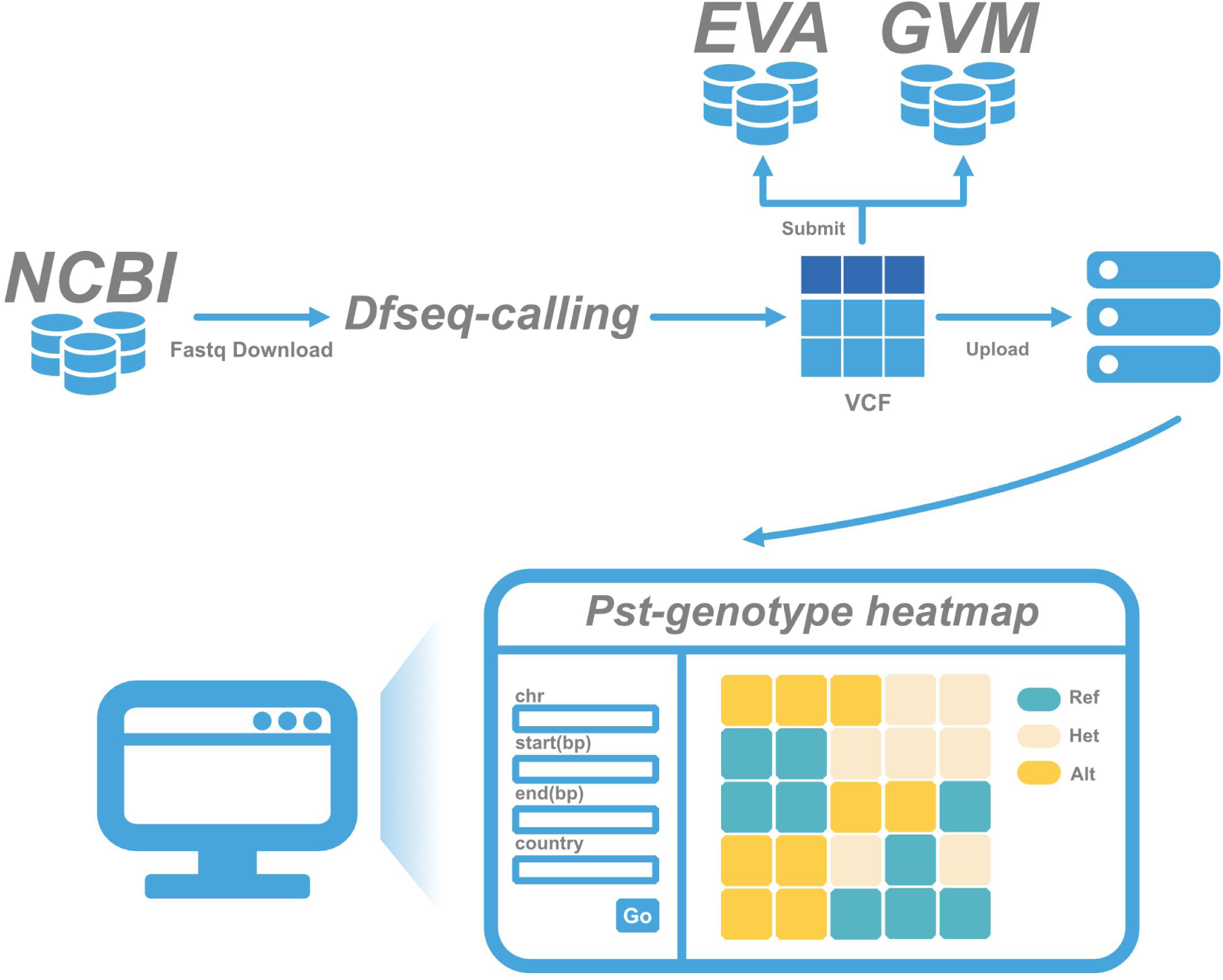
Construction of the global Pst genetic variation map and data accessibility. The construction of the *Pst* genetic variation map in this project. First, FASTQ files of worldwide *Pst* isolates were downloaded. Using the Dfseq-calling pipeline, genetic variants were detected, producing a comprehensive genetic variation map (VCF format). This VCF file was subsequently uploaded to the European Variation Archive (EVA), Genome Variation Map (GVM), and our own server. A platform was developed to visualize the global *Pst* genotypes, allowing users to retrieve genotypes for any genomic region. Additionally, the VCF file is available for download from the EVA and GVM databases for any future research.

**Fig.S9.**
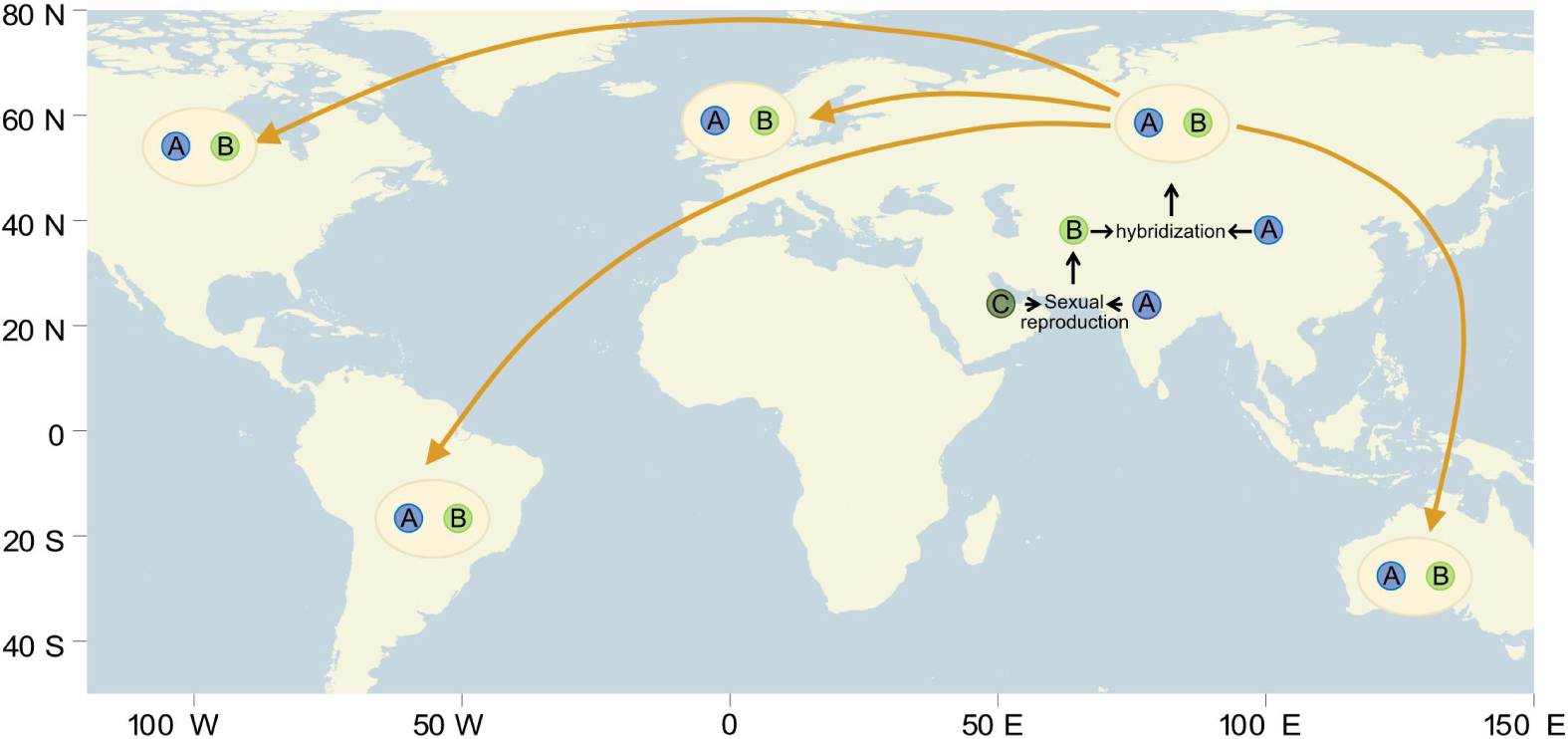
A hypothesis of the A/B haplotype origin. A hypothesis for the origin of the A and B haplotype. We suggest that their divergence resulted from one of the haplotypes undergoing a sexual recombination event with a hypothetical C haplotype. Afterward, the A and B haplotypes merged to form the AB haplotype isolate, which subsequently spread across the globe.

## Table

Table S1: Metadata of 266 global *Pst* isolates WGS public data Table S2: Alignment results of 239 *Pst* isolates

Table S3: Information of the genetic variation map, statistics using SnpEff Table S4: Genetic distance of 239 isolates to four haploid reference genomes

Table S5: Distribution of three different population variation landscapes regions in genome Table S6: Effector list

Table S7: Alignment results of 250 *Pst* transcriptome data Table S8: GO enrichment terms for mc-hpd and hc-hpd regions Table S9: Alignment results of 183 *M. oryzae* isolates

Table S10: Alignment results of 117 inbred population isolates Table S11: GWAS re-mapping results

Table S12: Phenotypic information of the inbred population

Table S13: Genome-wide distribution of recombination breakpoints in inbred population

## Acknowledgements

This work was supported by the Biological Breeding-Major Projects (2023ZD04076), the National Key Research and Development Program of China (2023YFF1000100), and the Research Program for Network Security and Information of the Chinese Academy of Sciences (CAS-WX2021SF-0109). The authors would like to thank J. Brown and M. McMullan for valuable suggestions.

## Declaration of Interests

The authors declare no competing interests.

## Author Contributions

Y.W. performed data analysis, developed the software and website, and wrote the first version of the manuscript. M.Y. contributed to data interpretation and manuscript writing. F.H. conceived the idea, coordinated data analysis and edited the manuscript.

## Data Availability

Genetic variation map of *Pst* have been deposited in the Genome Variation Map (GVM) in National Genomics Data Center, Beijing Institute of Genomics, Chinese Academy of Sciences and China National Center for Bioinformation, under accession number GVM000884 (https://ngdc.cncb.ac.cn/gvm/getProjectDetail?Project=GVM000884). It is also available at European Variation Archive, under accession number PRJEB82205 (https://www.ebi.ac.uk/eva/?eva-study=PRJEB82205).

## Notes

https://www.ebi.ac.uk/eva/?eva-study=PRJEB82205

https://ngdc.cncb.ac.cn/gvm/getProjectDetail?Project=GVM000884

### Competing Interest Statement

The authors have declared no competing interest.

